# Individual-versus group-optimality in the production of secreted bacterial compounds

**DOI:** 10.1101/094086

**Authors:** Konstanze T. Schiessl, Adin Ross-Gillespie, Daniel M. Cornforth, Michael Weigert, Colette Bigosch, Sam P. Brown, Martin Ackermann, Rolf Kümmerli

**Affiliations:** Department of Environmental Microbiology, Swiss Federal Institute of Aquatic Science and Technology (Eawag), Überlandstrasse 133, 8600 Dübendorf, Switzerland; Department of Environmental Systems Science, Swiss Federal Institute of Technology (ETH Zurich), Universitätsstrasse 16, 8092 Zürich, Switzerland; Department of Plant and Microbial Biology, University of Zürich, Winterthurerstrasse 190, 8057 Zürich, Switzerland; School of Biological Sciences, Georgia Institute of Technology, Atlanta, USA; Department of Health Sciences and Technology, Swiss Federal Institute of Technology (ETH Zurich), Universitätsstrasse 2, 8092 Zürich, Switzerland; Department of Quantitative Biomedicine, University of Zürich, Winterthurerstrasse 190, 8057 Zürich, Switzerland

**Keywords:** division of labor, bacteria, group level selection, siderophores, optimal production, economy of scales

## Abstract

How unicellular organisms optimize the production of compounds is a fundamental biological question. While it is typically thought that production is optimized at the individual-cell level, secreted compounds could also allow for optimization at the group level, leading to a division of labor where a subset of cells produces and shares the compound with everyone. Using mathematical modelling, we show that the evolution of such division of labor depends on the cost function of compound production. Specifically, for any trait with saturating benefits, linear costs promote the evolution of uniform production levels across cells. Conversely, production costs that diminish with higher output levels favor the evolution of specialization – especially when compound shareability is high. When experimentally testing these predictions with pyoverdine, a secreted iron-scavenging compound produced by *Pseudomonas aeruginosa*, we found linear costs and, consistent with our model, detected uniform pyoverdine production levels across cells. We conclude that for shared compounds with saturating benefits, the evolution of division of labor is facilitated by a diminishing cost function. More generally, we note that shifts in the level of selection from individuals to groups do not solely require cooperation, but critically depend on mechanistic factors, including the distribution of compound synthesis costs.

## Introduction

The production of proteins and metabolites is at the basis of many cellular processes. Because of its fundamental importance, there has been much interest in understanding the mechanisms that control compound production at the cellular level (Dekel and Alon 2005; Shachrai et al. 2010; Eames and Kortemme 2012; Price et al. 2013). The general consensus is that there should be a production optimum, at which the net benefit of producing a specific compound (protein or metabolite) peaks. Moving away from that optimum either results in wasteful overproduction or insufficient compound supply. Indeed, experimental work has demonstrated that populations of unicellular organisms evolve towards such optima in laboratory settings (Dekel and Alon 2005; Poelwijk et al. 2011).

One implicit assumption of the above-mentioned work on unicellular organisms is that optimality is achieved at the level of the individual. This means that every individual in a population is expected to tune its compound expression level to the same optimum. However, bacteria and other microbes are not solitary organisms, but often live in dense social groups, where they can interact and communicate with one another (West et al. 2007; Asfahl and Schuster 2017; Özkaya et al. 2017). This opens the possibility for optimality in compound production to arise at the level of the clonal group and not the individual. Indeed, recent experimental studies report cases of bimodal trait expression patterns across individuals in clonal groups, which could reflect group-level optimization (Diard et al. 2013; van Gestel et al. 2015).

Here, we formally ask whether group optimality in compound production in a clonal group is indeed possible and examine the conditions for which heterogenous suboptimal individual behavior can promote optimality at the level of the group (i.e. representing a case where clonal groups become a functional unit; (Ackermann 2015). (Frank 2013) proposed that heterogeneous expression profiles could be particularly common for secreted extra-cellular compounds. The reasoning is that secreted compounds, including enzymes and secondary metabolites, can be shared at the level of the clonal group, thereby decoupling individual expression levels from fitness returns. In other words, the benefit an individual gains from the production of a shared compound is determined not by the individual’s own investment, but by the production level of the entire group (Frank 2013). High shareability could thus potentially trigger the evolution of a group-optimal strategy, characterized by a type of division of labor where the clonal group splits into phenotypic producers and non-producers. This split could be mechanistically implemented either by intrinsic or extrinsic (signal-induced) bistable expression of the relevant synthesis genes (Dubnau and Losick 2006; Veening et al. 2008; Ackermann et al. 2008; West and Cooper 2016), combined with phenotypic switching promoting a stable proportion of producers and non-producers in the population (Julou et al. 2013; Diard et al. 2013).

Apart from the aspect of shareability, we further hypothesize that whether or not a shift from individual to group level optimization occurs also depends on the underlying cost and benefit functions of a compound. Previously published data show biological examples of costs that increase either linearly or exponentially with the amount of compound produced (Dekel and Alon 2005; Shachrai et al. 2010; Eames and Kortemme 2012; Price et al. 2013). In the case of a linear cost function, we predict that group-level optimization should not evolve because costs are additive, thus providing no fitness incentives for cells to engage in specialization. Conversely, diminishing cost functions should be more conducive for group-level optimization because the principle of the economies of scale apply – the cost per compound decreases with higher production levels (Case et al. 2011).

We mathematically formalized these hypotheses to define the conditions required for group-level optimization to evolve. We then empirically tested our hypotheses in a bacterial model system. Specifically, we used the production of pyoverdine, a siderophore synthesized via nonribosomal peptide synthetases, which is secreted by the bacterium *Pseudomonas aeruginosa* to scavenge iron from the environment (Visca et al. 2007). Pyoverdine production has become a model trait to study social interactions in bacteria, and multiple studies have shown that this compound can be shared between cells that possess the specific receptor required for pyoverdine uptake (e.g. (Griffin et al. 2004; Cordero et al. 2012; Inglis et al. 2016). However, the level of shareability for this and other compounds depends on a number of factors, including the diffusivity of the environment and cell density (Ross-Gillespie et al. 2009; Kümmerli et al. 2009a; Weigert and Kümmerli 2017). This means that the level to which pyoverdine can be shared as a public good among cells might vary in the environment and that is why we included shareability as a key parameter in our model.

Pyoverdine, like many other secreted secondary metabolites (e.g. antibiotics, toxins, surfactants, siderophores), is produced via nonribosomal peptide synthesis (NRPS) instead of ribosomal synthesis (Caboche et al. 2007; Miethke and Marahiel 2007). Nonribosomal peptide synthesis requires the buildup of complex synthesis machineries prior to metabolite production (at least 23 proteins in the case of pyoverdine; (Visca et al. 2007; Schalk and Guillon 2013). Given this mechanistic background, we postulate that if the cost of building up this dedicated cellular machinery is substantial compared to the production of the compound itself then the system should be characterized by a diminishing cost function. Conversely, if the majority of pyoverdine cost is attributable to molecule synthesis and not to the buildup of the biosynthetic machinery then the system should be characterized by a linear cost function. To test these predictions, we measured cost and benefit functions of pyoverdine production using an engineered *P. aeruginosa* strain, where pyoverdine synthesis can be gradually induced. Subsequently, we combined automated flow cytometry with single-cell microscopy to examine whether all individual cells within a clonal group show uniform pyoverdine production levels or whether specialization occurs. These analyses allow us to examine whether *P. aeruginosa* has been exposed to selection for optimal production at the group level in the evolutionary past, in its natural environment.

## Material and Methods

### Mathematical model

We used a simple mathematical model to explore how cost and benefit functions interact to shape bacterial fitness in a clonal group where individuals either all produce the same amount of compound (e.g. pyoverdine) or engage in specialization where only a fraction produces the compound. We can express the net benefit *β* to a bacterial group as

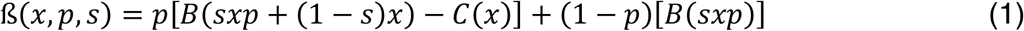

where *B* is a function representing the gross benefit of the compound, and *C* is the absolute cost of compound production. *p* is the proportion of producers in the group, producing a quantity *x* (the other individuals produce none), and *s* is the shareability parameter, describing the proportion of compound that is shared among clonemates. Our model focuses on a complete division of labor where production phenotypes are distributed in a binary on-off way. We chose this extreme setup for two reasons. First, biological examples of phenotypic division of labor include such binary switches (Ackermann et al. 2008; Diard et al. 2013; van Gestel et al. 2015). Second, although other scenarios of division of labor (e.g. where cells produce either low or high amounts of a compound) would also be realistic, we expect highest net group benefits for the on-off case. This is because with less extreme forms of specialization all cells would still experience some level of production costs, which would compromise the net benefits gained by specialization. We further assume that an individual’s production state (producer vs. non-producer) is non-heritable, and that individuals can change their state at any moment in time (see (Julou et al. 2013) for an empirical analysis of state switching). We note that the ability to switch production state is an important prerequisite for the evolution of phenotypic division of labor in a clonal group, as it guarantees a stable proportion of producers in the population.

The first term in the equation represents the fitness of a cell engaging in production of the secreted compound. The benefit experienced by a producer cell is a function of the amount of compound it can access, which is determined by the products released and shared by others (*sxp*) as well as those released by itself but not shared with others *(1-s)x*. Producers also pay a cost for production, which is a function of their production *(x)*. The second term describes the fitness of a non-producer, which equals the benefits obtained from others’ secretions.

We considered two monotonically increasing absolute cost functions: a diminishing cost function captured by *C*(*x*) = *g*(1-*e^−hx^*), and a linear cost function captured by *C*(*x*) = *k*(*x*), where *g, h*, and *k* are scaling factors (Figure 1A, B). Note that we use an arbitrary diminishing cost function for our main analysis, but provide a general mathematical proof that shows that our findings hold for any type of diminishing cost function (see Supplemental Information). For the gross benefit function, we assume saturating returns with higher compound availability, captured by *B*(*x*) = *d*(1-*e^−fx^*) with *d* and *f* as scaling factors (Figure 1A+B). Saturating returns are common for secreted metabolites (Foster 2004; Cornforth et al. 2012).

**Figure 1.**
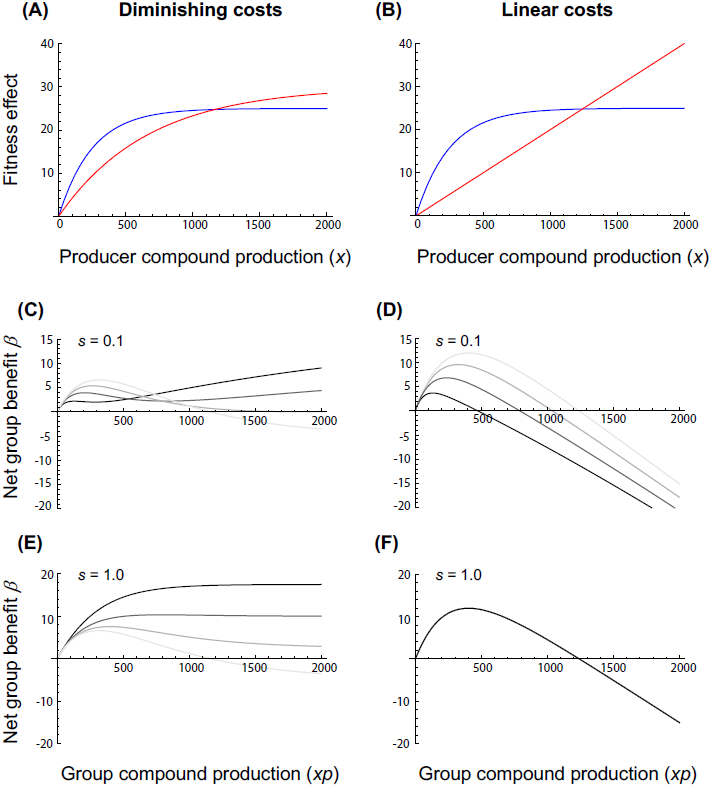
The net benefit to a group of bacteria producing a shared compound in relation to absolute compound production costs, shareability and the proportion of producers within the group. Panels A and B show examples of a diminishing and a linear absolute compound cost function *C* (red lines), respectively, in relation to a saturating gross benefit function *B* for producers (blue lines) assuming shareability s = 0. Panels C - F depict the resulting net benefit *β* = *B - C* to the group as a function of group compound production (*xp*), for different values of compound shareability s and degrees of production specialization *p* (grey shaded lines) among cells within the group: (C) diminishing costs and low shareability; (D) linear costs and low shareability; (E) diminishing costs and high shareability; (F) linear costs and high shareability. Grey shaded lines indicate scenarios for different proportions of producers, ranging from *p* = 1 (faint grey line), to *p* = 0.75 (light intermediate grey line), top = 0.5 (dark intermediate grey line), to *p* = 0.25 (dark grey line). Note that maximum net group benefits equal single-cell optima for *p* = 1.

### Strains

All our experiments were carried out with the clinical isolate *P. aeruginosa* PAO1 (ATCC 15692), the most widely studied strain of this species. To measure costs and benefits of pyoverdine production, we constructed the conditional pyoverdine mutant *PAO1-pvdSi*, where we put the production of pyoverdine under the control of the IPTG inducible promoter *P_tac_*. Specifically, we replaced the native promoter of *pvdS*, the gene coding for the iron starvation sigma factor PvdS, which controls the expression of genes coding for enzymes involved in nonribosomal peptide synthesis of siderophores. We PCR-amplified the flanking regions (FR1 upstream: 836 bp; and FR2 downstream: 987 bp) of the native *pvdS* promoter using the following set of primers (FR1 F: 5′-ggg ata cac cgg aga gga ata −3′, FR1R: 5′ - ttt gcc att ctc acc gga ttt gat gga ggg gag aaa ttc tt −3′ / *FR2F-SacI*: 5′ - gcg cga gct cat gtc gga aca act gtc tac ccg −3′, *FR2R-Kpnl*: 5′ – gcg cgg tac cat cca tat gcc cgc tgc gac a - 3′). FR2 features two restriction sites (*SacI* and *Kpnl*), which we used to clone this fragment into the pME6032 vector featuring the *P*_tac_ promoter and *lacI* (repressor of *P*_tac_). We then PCR-amplified the *lacI* - *P*_tac_ - FR2 construct using the primers ampF (5′-aat ccg gtg aga atg gca aa −3′) and ampR (5′-tcc gag gcc tcg aga tct at −3′), and subsequently carried out fusion PCR with this amplicon and FR1, which features a 20 bp overhang (see FR1R) reverse complementary to the *lacI* sequence. Fusion PCR included a first set of cycles, where the *lacI* - *P*_tac_ - FR2 amplicon and FR1 could hybridize via the overhang in the absence of primers, and therefore mutually serve each other as template and primer. During a second set of cycles, the final construct of 3242 bp length was amplified using the primers FR1 and ampR. This fragment was ligated into pGEM-T easy (A1360, Promega) for storage, and cloned into pME3087, a suicide vector, which we transformed into the *Escherichia coli* S17-1 donor strain. We then carried out mating between the S17-1 donor and the PAO1 recipient. Single crossovers were selected on LB agar supplemented with tetracycline (100 *μ*g/ml to select against wildtype PAO1) and chloramphenicol (10 *μ*g/ml to select against S17-1). We grew one single crossover colony overnight in LB at 37°C, during which double crossovers can occur (either reverting back to wildtype or replacing the native *pvdS* promoter with our inducible construct). To enrich for double crossovers, we added tetracycline (20 *μ*g/ml, a bacteriostatic concentration at which non-tetracycline resistant strains stop growing). Subsequently, we added carbenicillin (2000 *μ*g/ml) to selectively kill the growing tetracycline-resistant single crossovers. After cell washing and plating, we picked ~100 colonies from LB plates and screened for conditional pyoverdine mutants, using IPTG-induction assays and sequencing of the *pvdS* promoter site.

To investigate heterogeneity in pyoverdine expression, we used PAO1 *pvdA-gfp*, a transcriptional reporter (chromosomal insertion: *attB::pvdA-gfp*), which links transcription of *egfp* to the transcription of *pvdA*, one of the pyoverdine biosynthesis genes (Kaneko et al. 2007). As negative control (i.e. no GFP expression), we used the wildtype PAO1 strain obtained from Pierre Cornelis’ laboratory (Ghysels et al. 2004). As positive control, we used a constitutive GFP-expressing derivate of PAO1 (PAO1-gfp, chromosomal insertion: *att*Tn7:: P_tac_ - *gfp*) (Lambertsen et al. 2004). Finally, we generated the double-reporter strain PAO1 *-mcherry-pvdA-gfp* by integrating the constitutive promoter P_tac_ linked to mcherry (attTn7:: P_tac_ -*mcherry*) into PAO1 *pvdA-gfp* using the miniTn7 system (Choi and Schweizer 2006).

### Media and growth conditions

All experiments were carried out in casamino acids medium (CAA) containing 5 gl^-1^ casamino acids (Merck, Switzerland), 1.18 g l^−1^ K_2_HPO_4_*3H_2_O, 0.25 g l^−1^ MgSO_4_*7H_2_O, and 5.96 g HEPES (all from Sigma-Aldrich). To measure the cost of pyoverdine production, we grew our conditional pyoverdine mutant PAO1-*pvdSi* in CAA medium supplemented with 100 *μ*M FeCl_3_. Under these conditions, pyoverdine is not required for growth, and consequently we expect that IPTG-induced pyoverdine production will induce costs, which can result in a longer lag phase and/or a reduced growth rate of the bacterial culture. Thus, the cost *c* of pyoverdine production at a given induction level *i* can either be measured as *c* = *λ*_0_ - *λ*_i_ or *c* = *μ*_0_ - *μ*_i_, where *λ*_0_ and *μ*_0_ are the lag phase and the maximal growth rate of non-induced *PAO1-pvdSi* cultures, whereas *λ*_i_ and *μ*_i_ represent the lag phases and the maximal growth rates of induced *PAO1-pvdSi* cultures. We measured the *c* for 10 induction levels (IPTG in mM: 0.1, 0.11, 0.12, 0.13, 0.14, 0.15, 0.175, 0.2, 0.3, 0.5) in 15-fold biological replication using multimode plate readers (Tecan Infinite M200 PRO and Biotek Synergy MX). To control for possible adverse effects of IPTG itself, growth in the presence of IPTG was measured in both the *PAO1-pvdSi* strain (n=2) as well as in a pyoverdine knock-out mutant (PAO1 *ΔpvdD*) not containing an IPTG-inducible promoter (n=3; IPTG in mM 0, 0.1, 0.11, 0.12, 0.13, 0.14, 0.15, 0.175, 0.2, 0.3, 0.5). Induction experiments were carried out in 96-well plates, where we added approximately 10^5^ cells from an overnight LB culture to 200 *μ*l medium. Plates were incubated in the multimode plate reader at 37°C for 24 hours. Culture growth was measured as optical density (OD; absorbance at 600 nm) at 15 minute intervals. Cultures were shaken for 10 minutes prior to readings. Growth curves were fitted to OD trajectories from 0 to 16 hours of growth in R 3.1.1 (R Core Team 2015) using functions from the ‘grofit’ package (Kahm et al. 2010). The Richards growth model yielded the best fit for the observed growth trajectories, and was thus used to estimate *λ* and *μ* values for all treatments.

In addition to OD, we also quantified the production of pyoverdine over time for each IPTG induction level. Pyoverdine is a naturally fluorescent molecule, which can be quantified through excitation at 400 nm and emission at 460 nm (Kümmerli et al. 2009b). It has previously been shown that relative pyoverdine fluorescence correlates linearly with the absolute pyoverdine content in bacterial cultures (Dumas et al. 2013).

To measure the benefit of pyoverdine production, we grew *PAO1-pvdSi* in iron-limited CAA medium (using 100 mg ml^−1^ human-apo transferrin to bind iron and 20mM NaHCO3 as cofactor, both from Sigma-Aldrich, Switzerland), supplemented with various amounts of purified pyoverdine (pyoverdine concentration in *μ*M: 0, 5, 10, 25, 50, 100, 200, 400; in 6-fold biological replication each). Pyoverdine was extracted using the protocol described in (Dumas et al. 2013), and supplemented in four time intervals (after 2, 4, 6, and 8 hours of growth) to mimic the continuously increasing pyoverdine concentrations typically observed in growing cultures. Since *PAO1-pvdSi* was not induced with IPTG in these experiments, growth was entirely dependent on the supplemented pyoverdine. Accordingly, the benefit *b* of pyoverdine can be measured as *b* = *μ*_S_ - *μ*_0_, where *μ*_0_ and *μ*_S_ are the maximal growth rates of the unsupplemented and supplemented *PAO1-pvdSi* cultures, respectively. Measurement of growth kinetics and the analysis of growth curves were carried out as described above.

To quantify cell-to-cell variation in gene expression, we used a slightly modified version of the iron-limited CAA medium: 5 g Casein Hydrolysate (Merck), 0.25 g MgSO_4_*7H_2_O (Sigma-Aldrich), 0.9 g K_2_HPO_4_ (Sigma-Aldrich), and 5.96 g HEPES (Sigma-Aldrich). The pH was adjusted to 7 with 5M NaOH. After autoclaving, the medium was filtered through a 0.1 *μ*m-filter in order to remove particulate iron as well as background hindering the flow cytometry measurements (Cellulose Nitrate Membranes, 0.1 μm, 47 mm ø, Whatman). Iron availability was adjusted by addition of varying amounts of Fe(III)EDTA (Ethylenediaminetetraacetic acid Iron(III)-Sodium Hydrate, Sigma-Aldrich). In order to standardize EDTA concentrations across samples, iron-free EDTA (Ethylenediaminetetraacetic acid disodium salt dehydrate, Sigma-Aldrich) was added to reach a total EDTA concentration of 20 μM. Four iron limitation environments were tested: 0 μM, 1 μM, 5 μM and 20 μM Fe(III)EDTA. EDTA and Fe(III)EDTA were dissolved in nanopure water, filtered through a 0.1 μm filter, and added to CAA after autoclaving.

For flow cytometry and microscopy studies, the strains were streaked from a frozen stock on LB agar and incubated overnight. Single colonies were picked and inoculated in 5 ml LB and incubated at 37°C in an orbital shaker (220 rpm). After 16 hours of incubation, the culture was diluted 1:200 in 5 ml LB and regrown to the exponential phase, corresponding to an optical density of approximately 0.13. To start the experiment, all strains were diluted 1:2000 to reach a starting cell concentration of around 10^4^ cells/ml, and inoculated in 100 ml CAA. Cells were cultured in 100 ml-Schott bottles in a water bath at 37°C, oxygenated via stirring on a magnetic plate. The lid was removed and the bottles were covered with aluminum foil to allow the cultures access to oxygen. From these cultures, samples for flow cytometry and microscopy were taken.

### Automated flow cytometry

We used automated flow cytometry to monitor temporal changes in the distributions of *pvdA*-reporter-associated fluorescence in cultures of *P. aeruginosa* growing under different levels of iron limitation. Our setup, broadly similar to that of (Besmer et al. 2014), enabled us to quantify the GFP fluorescent signal in a time-resolved manner. To the intake port of a C6 flow cytometer (BD Accuri, San Jose CA, USA), we fitted an electronically-controlled relay valve that allowed switching between multiple sources. This setup permitted automated time-series sampling from up to 8 cultures in parallel. GFP-signal was detected using a 488 nm laser for excitation, and an emission detector fitted with a 533±30 nm bandpass filter. Forward- and side-scattered light produced by the 488 nm laser was also quantified, and all particles associated with a forward scatter signal of < 10000 fluorescence units were excluded *a priori* from further analysis on the basis that they were unlikely to represent intact bacterial cells. At each sampling point, the flow cytometer processed the sample at a rate of 66 μl min^−1^ for two minutes, although we retained data only from the final 20 seconds of each run, where the signal was most stable. Between each successive sample, the line was washed with 0.2% hypochlorite and nanopure water to avoid contamination and signal bleed-through. Cultures were sampled at 33-minute intervals during a total period of 180 minutes. The reporter strain, PAO1 *pvdA-gfp* was tested in four different Fe-limitation environments (CAA supplemented with 0 μM, 1 μM, 5 μM and 20 μM Fe(III)EDTA, respectively). We replicated the time-series experiment 8 times, where each block included the reporter strain in all four environments, a blank control (i.e. CAA medium only), a negative control for background GFP-fluorescence (i.e. PAO1 growing in unsupplemented CAA) and two positive controls (PAO1-gfp, under varying levels of iron limitation). Our raw flow cytometry data likely included noise – i.e. measurements from particles that were not single, intact bacterial cells. Therefore, for each sample at each time point, we calculated 95% contours of highest posterior density for the two-dimensional space of GFP-signal versus side scatter, and we excluded from further analyses any points lying outside these contours. These analyses, and all others described below, were performed in R 3.1.1 (R Core Team 2015).

### Microscopy

In order to collect single-cell data on pyoverdine expression, we acquired snapshots of bacterial cultures with an epifluorescence microscope. Bacteria were grown in conditions identical to the flow cytometry experiments, but only measured once every hour and only tested in CAA with 20 *μ*M EDTA, without addition of iron. At t=0h, cells were sampled directly from the LB medium. At 1, 2, and 3 hours cells were taken from the iron-limited medium and concentrated by centrifugation of 15 ml culture at 4°C, 5000 rpm, 10 minutes. Cells were kept on ice until the measurement and processed within 45 minutes.

For data acquisition, 1 *μ*l of cell suspension was spotted on a CAA 1.5% agarose pad and covered with a cavity slide as described in (Bergmiller et al. 2011). We used an inverted fluorescence microscope (Olympus IX81) with an X-Cite 120PV fluorescence lamp (Lumen Dynamics Group Inc., Canada) and an automated stage. Pictures were acquired with a cooled CCD camera (Olympus XM10) and an UPLFLN 100xO2PH/1.3 phase contrast oil immersion objective (Olympus). Phase contrast images were acquired with an exposure time of 20 ms, fluorescence images with an exposure time of 500 ms for gfp and 800 ms for mcherry. The filter for recording the GFP signal had the following parameters: excitation range 450-490, emission range 500-550 nm, DM 495 nm (AHF Analysentechnik, Tübingen, Germany; Article No. F46-002), the filter for mcherry: excitation range 540-580, emission range 595-665, DM 585 (AHF Analysentechnik, Tübingen, Germany; Article No. F46-008). Data for each type of experiment were recorded within one week in order to minimize variation of lamp intensity between experiments.

Growth potential of the cells was assessed by time-lapse microscopy. The growth of cells was followed by acquiring phase contrast and fluorescence pictures every 10 minutes for 4 hours, while cells were incubated at 37°C in a temperature-controlled setup (Cube and Box incubation system, Life Imaging Services, Reinach, Switzerland).

To follow microcolony growth on agarose pads, we used the same setup as described above, but this time inoculating pads with a highly diluted bacterial culture (dilution factor: 2.5*10^−4^). To induce iron limitation, we added 450 *μ*M 2,2′-bipyridin to the agarose pad. We then identified independent fields of view, which contained single cells (max. 5 cells), and followed the gene expression of all cells in the growing microcolony over time. Phase contrast and fluorescence pictures were acquired every 15 minutes for 5 hours. We obtained time-lapse data from 5 fields of views and gene expression data from approximately 1000 cells in total.

Analysis of microscopy pictures was conducted with ImageJ (freely available from http://rsbweb.nih.gov/ij/). First, the positions and outlines (‘masks’) of cells within images were determined from phase contrast pictures. Cells were automatically thresholded, converted into masks, and bigger cell groups were split apart with the watershed filter. The position of the cell masks was stored with the ‘analyzing particles’ command, with a size-based selection (250-Infinity in pixel units). These masks were then checked manually by overlaying over the phase contrast pictures, and all false positives were removed. The masks were then superimposed over the fluorescence images. The background was removed by subtracting a fluorescent image without cells, specific for the time point measured, from the fluorescence image. If no such image was available (i.e. for data for Figure 5), the average intensity of two cell-free regions was taken as background. Mean gfp-values were recorded for each cell, exported and analyzed in R 3.1.1. (R Core Team 2015).

## Results

### Modelling reveals that division of labor can only maximize group productivity if compound cost functions are diminishing and not linear

Our mathematical model investigated conditions for which the net benefit of compound production in a clonal group of cells is maximized by uniform production or division of labor, respectively. Specifically, we modelled the influence of the cost function (diminishing costs Fig. 1A; linear costs Fig. 1B) and shareability on the net fitness *ß* at various proportions of producer cells in a group. In the case of a diminishing cost function and low levels of compound sharing, our model reveals that whether *ß* is maximized via specialization or uniform production levels depends on the compound production level (Figure 1C). Specifically, low compound production maximizes the net benefit *ß* for uniform production levels across cells because compound concentration is simply too low to benefit everyone in the group. Conversely, high compound production levels guarantee that still a significant amount of compounds reaches nonproducing neighboring cells, despite low shareability. When considering high levels of compound sharing, our model reveals that *ß* is always maximized through specialization (Figure 1E). Thus, our model suggests that with diminishing costs and high shareability we would expect bacteria to evolve molecular mechanisms to restrict compound production to a fraction of the individuals in a clonal group.

In contrast, when considering linear cost functions we found that specialization had either a negative (Figure 1D, low shareability) or no effect on *ß* (Figure 1F, high shareability). For linear costs, we thus expect selection to favor a uniform level of compound production across cells within a clonal population. In line with previous findings (Dekel and Alon 2005; Poelwijk et al. 2011), in all cases there is one optimal production level that maximizes the net benefit of groups.

### Pyoverdine production entails linear costs and generates saturating returns

To estimate the cost function of pyoverdine, we induced its production in our engineered conditional mutant (PAO1-*pvdSi*) across a range of IPTG concentrations in iron-replete media. In this media, pyoverdine is not needed and the induction of its synthesis should generate costs, possibly manifesting in an extended lag phase and/or a reduced growth rate. Compatible with these expectations, we found that increased levels of pyoverdine induction led to both longer lag phases and lower growth rates (Figure 2A, B). These costs did not manifest when a non-inducible strain was exposed to IPTG, showing that the observed costs in PAO1-*pvdSi* are attributable to pyoverdine production and not to potential adverse effects of IPTG (Figure S1).

**Figure 2.**
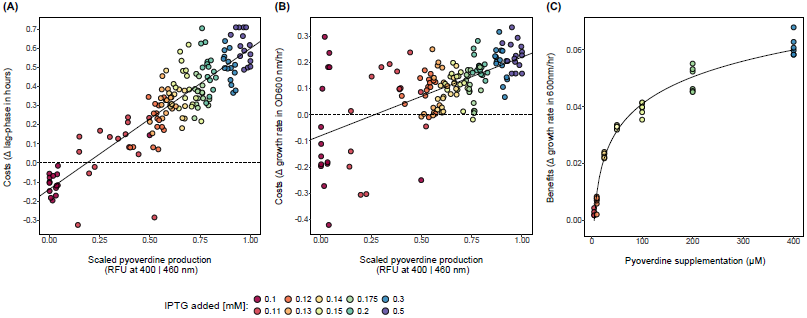
Cost and benefit functions of pyoverdine production. Costs were measured by inducing pyoverdine production in strain PAO1-*pvdSi* by the addition of increasing concentrations of IPTG under conditions where it is not required for growth. (A) shows costs detectable as increased lag phase, and (B) shows costs detectable as decreased growth rate. The pyoverdine peak value was normalized to the pyoverdine peak at maximum induction (0.5 mM IPTG). For both lag phase and growth rate costs, a linear function was the best fit, as indicated in the graphs (lag-phase: *F*_1,146_ = 499.5, *R*^2^ = 0.772, *p* < 0.0001; growth rate: *F*_1,146_ = 90.9, *R*^2^ = 0.379, *p* < 0.0001). To quantify benefits, purified pyoverdine was added to cultures of non-induced *PAO1-pvdSi* under iron-depleted conditions (C). We measured the increase in maximal growth rate (relative to unsupplemented conditions) as a function of pyoverdine supplementation. Here a logarithmic function (y = 0.01372 * log(x) −0.02216, AIC = −345.4) explained more variance than either a linear (AIC = −252.8), quadratic (AIC = −285.7), or an exponential (AIC = −299.1) fit. Cost and benefit functions cannot directly be overlaid because the assays occurred in CAA media with different iron concentrations, affecting the absolute growth rate and fitness measurements (on the y-axis). However, the x-axis range is comparable for both assays, assuming that 0.5 mM IPTG induces nearly maximal pyoverdine production and knowing that *P. aeruginosa* maximally produces around 300 μM pyoverdine in strongly iron-limited CAA over 24 hours (Dumas et al. 2013).

When fitting statistical models to the cost data, we found that both lag phase extensions and growth rate reductions scaled linearly with induced pyoverdine production levels (Figure 2A, B). Interestingly, we found that a low induction of PvdS actually resulted in a net benefit (i.e. negative costs) mainly in terms of growth rate under iron-replete conditions. This finding is compatible with the view that PvdS as a sigma factor regulates, in addition to pyoverdine production, a number of other genes (Tiburzi et al. 2008), such that low PvdS expression is beneficial even when pyoverdine is not needed.

Finally, we estimated the benefit function of pyoverdine by supplementing PAO1-pvdSi with different concentrations of purified pyoverdine in iron deplete media (Figure 2C). We found that benefits saturated with increasing pyoverdine concentrations, indicating that above a certain level of pyoverdine availability iron is either no longer the limiting resource or its uptake can no further be increased.

### Gene expression analyses indicate uniform pyoverdine production across cells

Our modeling insights combined with the discovered linear cost function for pyoverdine production would predict uniform pyoverdine expression levels across individuals in a clonal group. To test this hypothesis, we quantified the expression of *pvdA* (encoding a pyoverdine synthesis protein) using the transcriptional reporter strain PAO1 *pvdA-gfp*.

In a first analysis, we measured GFP expression of individual cells in a time-resolved manner using automated flow cytometry in environments differing in iron concentration. Iron availability alters the costs and benefits of pyoverdine production (Kümmerli et al. 2009b), which could potentially impact both individual- and group-level investment profiles. The data indeed show fine-tuned adjustments of mean pyoverdine expression levels in response to iron limitation and time (Figure 3A), indicating that *P. aeruginosa* adjusts pyoverdine expression to match prevailing levels of iron limitation. When looking at the individual-cell level, the distribution of GFP-signal intensity followed a unimodal distribution, with considerable variation around the mean but no signs of bimodality in *pvdA* expression across cells (Figure 3B).

**Figure 3.**
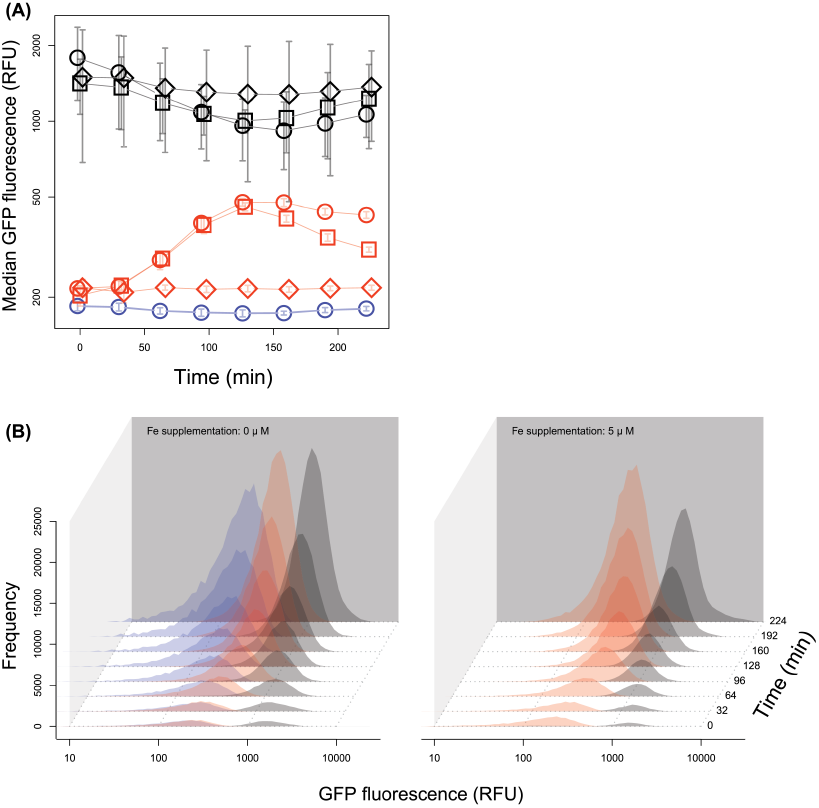
Tuning of pyoverdine expression in response to iron availability and time. (A) Population-level median GFP expression over time for the *pvdA-gfp* reporter (red lines) and a control strain constitutively expressing GFP (grey lines) under different iron supplementation regimes (circles = 0 *μ*M iron added, squares = 5 *μ*M iron added, diamonds = 20 *μ*M iron added). The blue line depicts background fluorescence signal of the wildtype PAO1 strain without reporter. (B) shows the distributions of individual-level GFP expression of the *pvdA-gfp* reporter (red), the constitutive *gfp* reporter (grey), and the wildtype control (blue) for the 0 *μ*M and 5 *μ*M iron supplementation regimes, respectively. While the data indicate that the absolute level of pyoverdine expression is extremely fine-tuned in response to iron availability and over time, in all cases the observed values are clearly unimodally distributed.

We then assessed inter-individual variation in *pvdA* expression using single-cell microscopy, a method that yields higher resolution of gene expression differences, especially at low expression levels. In a first experiment, we analyzed the *pvdA* expression of single cells grown under well-mixed conditions, i.e. the same conditions used for the flow cytometry experiment. This analysis confirmed our observation that *pvdA* expression followed a unimodal and not a bimodal pattern at early time points of the experiment (Figure 4A). At later time points we observed the emergence of a subpopulation of individuals that did not express *pvdA*, but these cells neither grew nor expressed a second constitutively expressed control fluorophore (mCherry; Figure 4B, C). This suggests that the cells not producing pyoverdine were generally inactive, a status possibly induced by the substrate shift from rich to minimal medium (van Heerden et al. 2014; Kotte et al. 2014).

**Figure 4.**
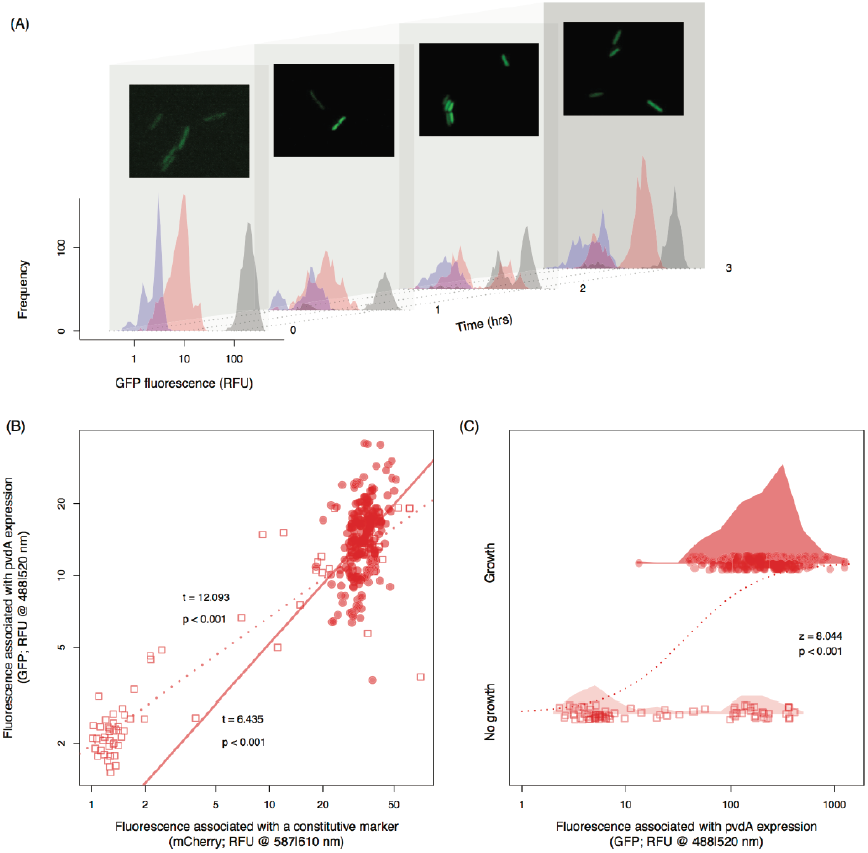
Microscopy confirms unimodal *pvdA* expression among growing cells. (A) Histograms show distributions of individual-level GFP expression of the *pvdA-gfp* reporter (red), the constitutive *gfp* reporter (grey), and the wildtype strain without *gfp* reporter (blue) over time. Microscopy pictures show representative snapshots of the PAO1 *pvdA-gfp* strain (brightness and contrast were adjusted manually). Data reveal a significant increase in bimodality in pyoverdine expression over time (linear increase of Hartigan’s *Dn*, diptest for bimodality (Maechler and Ringach 2013), for the *pvdA-gfp* reporter: *t*_66_ = 2.85, p = 0.006, but neither for the constitutive *gfp* reporter control: t_66_ = 1.16, p = 0.249, nor for the wildtype strain without *gfp*: t_66_ = −0.94, p = 0.352). However, (B) fluorescence from expression of a constitutive *mcherry* control marker served as a significant linear predictor of fluorescence from expression of pyoverdine (*pvdA-gfp*) both for growing (filled circles) and non-growing cells (empty squares). This suggests that pyoverdine expression levels are linked to the overall cellular gene expression activity. (C) Bimodality in *pvdA-gfp* expression is only observed among non-growing cells (squares), but was absent among dividing cells (circles; ≥ 2 divisions observed within four hours) (*Dn* = 0.024, p = 0.485). This indicates that bimodality in pyoverdine expression at the whole population level is mainly driven by bistability in the growth status of cells (dashed line: significant logistic regression between a cell’s pyoverdine (*pvdA-gfp*) expression and its growth status). Shaded areas show density functions of *pvdA-gfp* expression levels.

An implicit assumption in the above experiments is that bimodality, had it indeed been observed, would have been facilitated at a mechanistic level simply by intrinsic noise in expression of the genes making up the regulatory network for our trait of interest. However, it could also be that bimodal expression of compound production would be more clearly seen if, in addition to the intrinsic gene expression noise, there was additional variability in the extracellular stimuli to which the gene regulatory network is responding. Specifically, we postulated that the overall range of possible pyoverdine expression levels could be widened if we introduced spatial heterogeneity in local pyoverdine concentrations by creating spatially structured environments. To assess this possibility, we performed a second microscopy experiment, where we grew micro-colonies of cells on agarose pads and followed the *pvdA* expression of individual cells over time (similar to (Julou et al. 2013). This experiment also revealed a unimodal and not a bimodal pyoverdine expression pattern (Figure 5).

**Figure 5.**
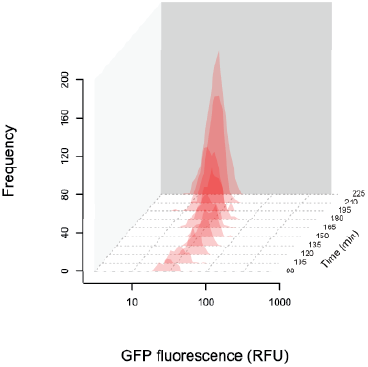
The expression of *pvdA* is also unimodal when cells grow in a spatially structured environment. Histograms show distributions of individual-level GFP expression of the *pvdA-gfp* reporter (red) over time. Cells were inoculated at low density onto agarose pads and their growth and individual *pvdA* expression was tracked over time using automated time-lapse microscopy. Data show gene expression profiles in 30-minutes intervals, starting from 90 minutes after the start of the experiment.

Taken together, our flow cytometry and microscopy data are consistent with the perspective that optimality in pyoverdine production in the natural environment is selected for at the level of the individual cell and not the group.

## Discussion

(Frank 2013) proposed that phenotypic specialization within clonal groups of microbes could be common for traits entailing the secretion and sharing of beneficial extra-cellular compounds between cells. Here, we formally addressed this hypothesis by combining mathematical modelling with empirical experiments. In line with Frank’s (2013) proposition we found that high shareability of secreted compounds is important for division of labor to evolve. However, in our model, high shareability was neither sufficient nor the key driver of specialization. Instead, our model showed that whether or not the evolution of single-cell or group-level optima is favored depended on the cost function of the trait. If costs scale linearly with production levels then uniform production levels maximize net benefits, whereas in case of diminishing costs division of labor does (Figure 1). When testing these predictions for pyoverdine, a secreted and shareable iron-scavenging molecule produced by *P. aeruginosa*, we found that costs scaled linearly with production levels. When probing whether division of labor is present for pyoverdine production, we detected unimodal and not bimodal gene expression pattern across cells. This finding is in agreement with our model and suggests that linear production costs combined with the environmental conditions this species faces in natural settings has selected for pyoverdine investment optimization at the individual rather than the group level.

We chose pyoverdine as a model trait because it features many of the putative prerequisites required for specialization to occur. At the gene regulatory level, it is well established that feedback loops can generate heterogeneity in gene expression (Alon 2007; Ackermann 2015). Compatible with this prerequisite, pyoverdine synthesis is regulated by two interacting feedback loops. There is a positive feedback loop up-regulating pyoverdine synthesis through a cell membrane embedded receptor-anti-sigma-factor signaling cascade (Lamont et al. 2002), and a negative feedback loop operating via Fur (ferric uptake regulator, (Ochsner and Vasil 1996). At the production level, pyoverdine is synthesized via NRPS, requiring the buildup of a large assembly line before compounds can be produced. These initial costs (known as standing costs in economy) could promote the economies of scale: a few specialists produce large quantities of a product, which distributes the initial fixed costs over many units of output (Case et al. 2011). Finally, we know that pyoverdine (and siderophores in general) can be shared under many conditions, including laboratory (Weigert and Kümmerli 2017) and natural (Cordero et al. 2012; Butaitė et al. 2017) settings.

Given these prerequisites, we might ask why cells showed uniform investment levels in our system, and what can we extrapolate from our findings to other microbial traits? For one thing, our results show that NRPS does not automatically lead to a diminishing cost function. Although perhaps counterintuitive at first, our findings match recent reports that the pyoverdine production machinery is metabolically inexpensive, only making up one hundredth of the biomass that flows into the production of the pyoverdine molecules themselves (Sexton and Schuster 2017). Although this estimate might vary in response to nutrient availability (Brockhurst et al. 2008; Xavier et al. 2011; Sexton and Schuster 2017), it highlights that secondary metabolite production via NRPS or polyketides (Donadio et al. 2007; Caboche et al. 2007; Miethke and Marahiel 2007; Weissman and Müller 2008) might often follow linear cost patterns. Thus, diminishing cost functions might be rare even for secreted and shareable compounds.

Moreover, even in a case where costs are diminishing, our model shows that the evolution of division of labor also critically relies on high shareability of benefits. This is in line with previous models showing that the degree of compound sharing strongly influences what type of social interaction is favored by selection (Estrela et al. 2016). Although high shareability of pyoverdine (and other secreted compounds) was demonstrated in controlled experiments (e.g. *s* = 0.95 in well-shaken cultures; (Inglis et al. 2016), several factors could contribute to reduced sharing in natural settings, thereby compromising the efficiency of specialization (Oliveira et al. 2014). First, in certain environments including aquatic systems bacterial cells are typically dispersed and occur at low density (Whitman et al. 1998), conditions that have been shown to reduce compound sharing (Ross-Gillespie et al. 2009; Scholz and Greenberg 2015). Second, compound shareability is determined by the diffusivity of the environment (Driscoll and Pepper 2010; Allen et al. 2013; Frank 2013). High diffusivity might reduce compound sharing between neighboring individuals as compounds rapidly diffuse outside the range of cells (Kümmerli et al. 2014). Similarly, low diffusivity (e.g. in environments that are not water saturated) likely reduces shareability because compounds stay close to producers (Kümmerli et al. 2009a; Julou et al. 2013; Weigert and Kümmerli 2017). Finally, bacterial groups might often not be clonal, in contrast to our model assumptions. Instead, producers could be surrounded by wholly unrelated microbes that neither partake in compound production nor uptake, but simply represent physical obstacles, thereby impeding efficient metabolite exchange among clonemates (Inglis et al. 2016). Alternatively, the community could include exploitative genotypes that use but never contribute to compound production, either within the same species (Griffin et al. 2004; Cordero et al. 2012) or across species (Cordero et al. 2012; Butaitė et al. 2017). In the presence of exploitative genotypes, producers of siderophores were observed to increase their production to high levels (Kümmerli et al. 2009b), presumably leaving little room for intra-clonal division of labor to evolve.

While our model focused on the role of compound production costs and shareability, there are clearly further biological and physical factors that affect the evolution of division of labor. For example, cell aggregation or other factors that promote group cohesion have been identified as essential elements driving the selection for division of labor (Wakano et al. 2009; Rossetti et al. 2010; Ratcliff et al. 2012b; Olejarz and Nowak 2014). More generally, evolutionary theory suggests that division of labor is difficult to evolve in loose groups of microbes, where the identity and frequency of partners can change rapidly and conflicts among different genotypes might be prevalent (West et al. 2015; West and Cooper 2016). This might especially hold true in fluctuating environment, where conditions favoring and opposing the evolution of division of labor might quickly alternate.

Despite these constraints, a number of studies showed that a genetic form of division of labor can evolve in laboratory settings (Ratcliff et al. 2012a; Kim et al. 2016; D′Souza and Kost 2016; Dragoš et al. 2018). One issue with genetic forms of division of labor is that the frequency of the two interacting partners required for group optimality might not be stable. This means that in the absence of any mechanism stabilizing strain frequency, one cell type could become dominant and thereby lower group performance (Tasoff et al. 2015; Kallus et al. 2017; Taillefumier et al. 2017). The above scenarios are different from the one considered in our study, where division of labor can occur in clonal populations through phenotypic specialization, and where phenotypic switching between compound production and non-production could guarantee a stable proportion of the two phenotypes. This form of division of labor has been reported for a number of systems (infection: (Ackermann et al. 2008; Diard et al. 2013); metabolism: (Flores and Herrero 2009); motility: (van Gestel et al. 2015). However, unlike these examples where division of labor is explained by trade-offs and task incompatibility, our theoretical model predicts that diminishing production costs of a single shareable compound could drive selection of phenotypic division of labor. While pyoverdine production did not allow us to test this hypothesis directly, due to its linear costs, we hope that our novel hypothesis will stimulate future research in the field, and result in the testing of our predictions across a wider range of social traits.

## Acknowledgements

We thank Frederik Hammes for support with flow cytometry. KTS, CB, and MA were supported by Eawag; KTS and MA were supported by ETH Zurich; ARG, MW and RK were supported by the Swiss National Science foundation (grant no. PP00P3-139164) and the European Research Council (grant no. 681295); KTS was supported by the Swiss National Science foundation (grant no. PP2EZP3-162260); MA was supported by the Swiss National Science foundation (grant no. 31003A_149267); MW was supported by DAAG, and DMC was supported by a University of Edinburgh studentship. The authors declare no conflict of interest.

