## supplemental information for "Individual-versus group-optimality in the production of secreted bacterial compounds"

### Supplementary Information

#### Supplementary modelling

##### Division of labor is optimal for any decelerating cost function if shareability is sufficiently high

Though the main text assumes specific cost and benefit functions, the advantage of division of labor holds in the model from the main text as long as the cost function is decelerating and shareability is sufficiently high.

**Theorem 1: Assuming the fitness function in the main text, for any differentiable benefit and cost functions (B(x) and C(x) respectively), with $x \in [0, \infty)$, that cross through the origin [$C(0) = B(0) = 0$] where the cost function has negative concavity [$C''(x) < 0$ for all x], there exists a value $0 \leq s^* < 1$ such that for any shareability constant $s > s^*$, fitness decreases as *p* decreases if *xp* is held constant (or equivalently, fitness increases as *x* increases, if *xp* is constant). In other words, assuming total production ($xp$) is constant, at a sufficiently high shareability level, fitness increases as the proportion of non-producers does.**

Proof:

First we show that the fitness function from the main text increases with *x* for the case where $s = 1$. If we fix $xp$ (letting constant $q = xp$), we can express the fitness function as

$$\beta_2(x,s) = \frac{q}{x}\left(B(qs + (1-s)x) - C(x)\right) + \left(1 - \frac{q}{x}\right)B(qs) \qquad (2)$$

The derivative of this function with respect to *x* ($\frac{d}{dx}\beta_2(x,s)$) simplifies to $p\left(C(x) - xC'(x)\right)/x$ for the case where $s = 1$. Because $p/x$ is positive, if $C(x) - xC'(x)$ is positive for all *x*, then the whole derivative is positive. Since $C(x)$ has negative concavity and goes through the origin, the line tangent to $C(x)$ at any fixed $x_1$ crosses the x-axis to the left of the origin. Call this tangent line $L(x) = xC'(x_1) + b$ (where $b > 0$). So $L(x_1) = C(x_1) = xC'(x_1) + b$. Therefore $C(x_1) > xC'(x_1)$, and more generally $C(x) > xC'(x)$ over the interval.

This means when $s = 1$, $\frac{d}{dx}\beta_2(x,s) > 0$. Because $\frac{d}{dx}\beta_2(x,s)$ is continuous, there exists a $0 \leq s^* < 1$ such that for all $s \in [s^*, 1]$, $\frac{d}{dx}\beta_2(x,s) > 0$. Or, in words, when the shareability constant (*s*) is sufficiently high (with no restrictions on *p* or *x*), fitness is maximized at arbitrarily low *p* and high *x*. ∎

**Division of labor is not generally optimal for a decelerating cost function if shareability is sufficiently low**

On the other hand, if public goods are *not* freely shared with others, then division of labor is not generally beneficial. At an extreme, when there is no sharing ($s = 0$), then $\beta(x,p,0) = p[B(x) - C(x)]$; and so fitness is maximized at $p = 1$ with some corresponding *x*. With an additional assumption that $C(x) - xC'(x) > B(x) -$

$xB'(x)$ in some interval $[a, b]$ then the maximal fitness for a given *xp* value (with $x$ in the interval $[a, b]$) occurs at *p*=1 for any shareability level below a cutoff.

**Theorem 2: Suppose there are differentiable benefit and cost functions (B(x) and C(x) respectively), with $x \in [0, \infty)$, that cross through the origin [$C(0) = B(0) = 0$]. If $C(x) - xC'(x) > B(x) - xB'(x)$ over some interval $[a, b]$ then there exists a value $0 \leq s^* < 1$ such that for any shareability constant $s < s^*$, the maximal fitness for a given *xp* (with *xp* in the interval $[a, b]$) occurs at *p*=1.**

The derivative of the function $\beta_2$ with respect to *x* ($\frac{d}{dx}\beta_2(x, s)$) simplifies to $p\left[\left(xB'(x) - B(x)\right) - \left(xC'(x) - C(x)\right)\right]/x$ for the case where $s = 0$.

Because $p/x$ is positive and $\left(xB'(x) - B(x)\right) - \left(xC'(x) - C(x)\right)$ is negative in the interval $[a, b]$, then the whole derivative is negative. This means when $s = 0$, $\frac{d}{dx}\beta_2(x, s) < 0$ in this interval. Because $\frac{d}{dx}\beta_2(x, s)$ is continuous, there exists a $0 \leq s^* < 1$ such that for all $s \in [0, s^*]$, $\frac{d}{dx}\beta_2(x, s) < 0$ in the interval $[a, b]$. So when the shareability constant (*s*) is sufficiently low, for a given value of *xp*, fitness is maximized at *p*=1.

In the example in the main text, $C(x) - xC'(x) > B(x) - xB'(x)$ for *x* values high enough to optimize fitness, so there is typically a threshold below which heterogeneity is disfavored in this example.

**Supplementary Figure 1**

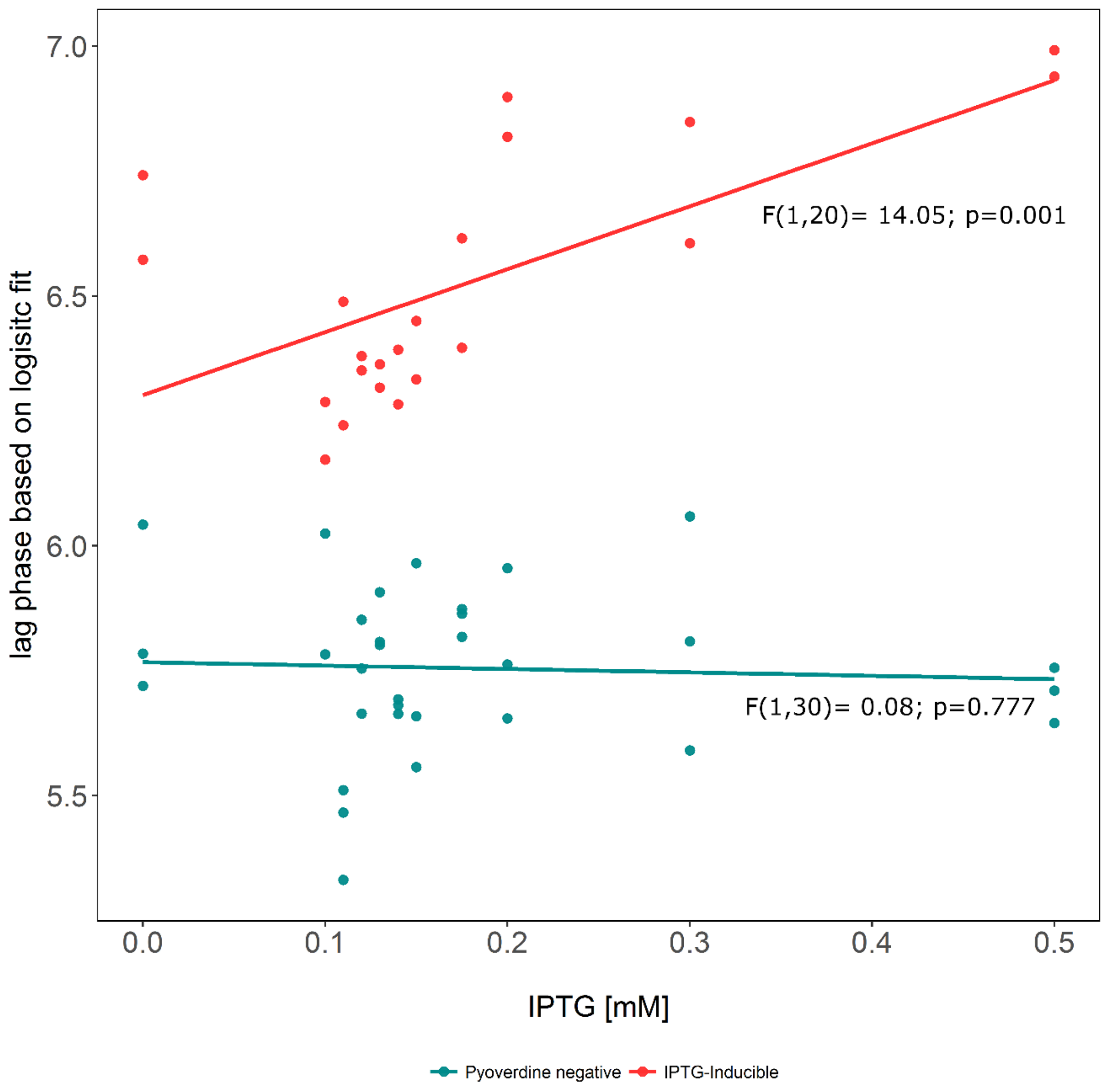


**Addition of IPTG itself has no effect on the lag phase.** Costs, visible as an increase in lag phase in a PAO1-*pvdSi* strain in the presence of IPTG (red; n=2) are caused by pyoverdine production and are not a side effect of IPTG, since the lag phase of a strain without IPTG-inducible promoter (PAO1*ΔpvdD*; blue; n=3) is not affected. Growth was monitored in a medium where pyoverdine is not required.

IPTG concentrations were varied (IPTG in mM 0, 0.1, 0.11, 0.12, 0.13, 0.14, 0.15, 0.175, 0.2, 0.3, 0.5) and the resulting growth curve was measured in a plate reader. We deducted the Richards Fit and calculated the lag phase (plotted on the y-axis). Using a linear fit, we find that the IPTG concentration significantly influences the lag phase in PAO1-*pvdSi*, but not in PAO1*ΔpvdD* (F-test results indicated in graph).
